# Tractor Workflow Pipeline: A Scalable Nextflow Framework for Local Ancestry-Aware Genome-Wide Association Studies

**DOI:** 10.1101/2025.09.02.673402

**Authors:** Nirav N. Shah, Taotao Tan, Jessica Honorato-Mauer, Yi-Sian Lin, Adam X. Maihofer, Clement C. Zai, PGC-PTSD ancestry working group, Marcos Santoro, Caroline M. Nievergelt, Elizabeth G. Atkinson

## Abstract

The routine exclusion of admixed individuals from traditional Genome-Wide Association Studies (GWAS) due to concerns about spurious associations has hindered genetic analyses involving multiple ancestries. *Tractor* GWAS addresses this issue by incorporating local ancestry into its analysis, empowering identification of ancestry-enriched hits and generating ancestry-specific summary statistics. However, *Tractor* requires accurate genomic phasing and local ancestry inference as prerequisite steps, which requires additional bioinformatics expertise and decision points regarding reference panel setup. To streamline, harmonize, and automate this process, we present a scalable *Nextflow* workflow that integrates all necessary steps, minimizing the need for manual intervention while remaining modular and customizable. The workflow supports multiple commonly used tools and offers flexibility in how *Tractor* is implemented. To demonstrate its utility, we applied this pipeline to analyze 32 blood biomarkers in 6,245 two-way AFR-EUR admixed individuals from the UK Biobank. This pipeline ran efficiently at scale, replicated known associations, and identified novel ancestry-specific loci. These novel associations were largely driven by variants present on African ancestral tracts but absent from European tracts, underscoring the value of local ancestry-aware methods in uncovering previously missed genetic signals. By enabling the efficient analysis of admixed individuals, our workflow facilitates *Tractor* use, paving the way for more broader genetic discovery.

## Introduction

Genome-Wide Association Studies (GWAS) have revolutionized the landscape of genetic research by identifying thousands of loci associated with complex traits and diseases, thereby deepening our understanding of their genetic architecture and informing the discovery of therapeutic targets (Sklar *et al*., 2011; Ikeda *et al*., 2018; Pardiñas *et al*., 2018; Ripke *et al*., 2014). Despite this progress, GWAS has traditionally excluded individuals with admixed individuals due to statistical challenges (Tan and Atkinson, 2023), limiting the generalizability of GWAS findings (Popejoy, Alice B., Fullerton, 2016; Sirugo *et al*., 2019; Mills and Rahal, 2020; Martin *et al*., 2019). While recent studies have begun to increasingly include individuals from multiple ancestry groups in meta- or mega-analysis approaches, participants with mixed ancestry are still often excluded in early filtering stages if they do not classify well to a particular ancestry assignment. To address this issue and improve analyses of admixed populations, our research group has developed *Tractor*, a local ancestry informed GWAS approach that is able to perform well-calibrated regression on admixed individuals alone or as part of cohorts with different ancestries (Atkinson *et al*., 2021). Local ancestry, which refers to the ancestral origin of specific chromosomal segments within an individual’s genome, provides a finer-scale resolution of genetic background than global ancestry estimates. *Tractor* tackles the known statistical challenge of differing genetic architecture across populations by taking local ancestry into account, allowing for fine-scale control of allele frequency stratification at the haplotype level.

While *Tractor* has been a foundational resource to facilitate GWAS in admixed populations, running *Tractor* involves several complex steps, creating a barrier to entry for groups with more limited in-house bioinformatics training or population genetics expertise. Specifically, since *Tractor* factors local ancestry estimates into the regression, we must phase data and determine the ancestry of haplotypic segments for each chromosome prior to regression, both of which are informed by a well-curated reference appropriate for the target population. The *Tractor* pipeline itself is also composed of several scripts for data pre-processing and subsequently GWAS. The Tractor pipeline consists of multiple scripts for data preprocessing and subsequent GWAS with several steps and decision points such as selecting optimal reference panels and linking intermediate results between stages. Automating and streamlining the entire workflow from genotype data through *Tractor* GWAS using pre-curated and benchmarked resources reduces manual effort and ensures reproducibility. This integration will facilitate broader use of this foundational tool and promote the inclusion of admixed samples in genetic studies.

Here, we present a Nextflow Workflow, a user-friendly and comprehensive pipeline that allows for running *Tractor* seamlessly across distinct compute environments (Di Tommaso *et al*., 2017). Nextflow is a Workflow Development Language used to create scalable, portable, and reproducible workflows. Its operability across local computers, High-Performance Computing (HPC) schedulers such as Slurm, AWS, Azure, and Google Cloud, makes it highly versatile. Additionally, it supports multiple ways to manage software dependencies, including Conda, Docker, and Singularity. This makes the *Tractor*Nextflow pipeline highly portable, requiring only basic pre-setup and configurations for the compute environment by the end-user.

Our Nextflow Workflow is divided into three independent modules: (1) Haplotype Phasing, (2) Local Ancestry Inference (LAI), and (3) Tract Extraction and *Tractor* GWAS (Fig. 1). This modular design enables users to customize the pipeline to suit their specific needs either by running all modules sequentially as part of the complete workflow or by substituting individual modules to incorporate alternative tools not currently supported by the pipeline.

**Fig. 1:**
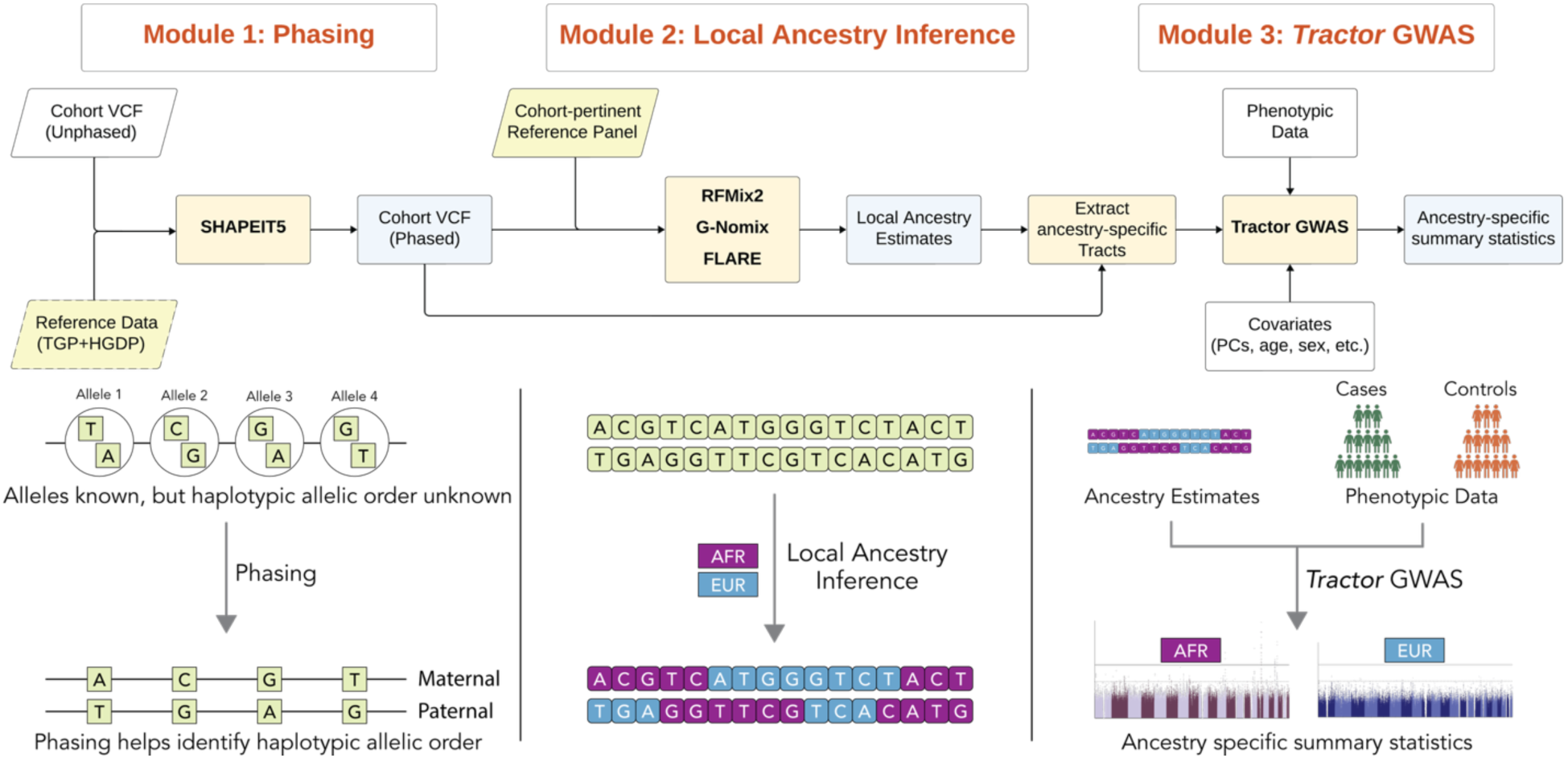
Overview of *Tractor* Nextflow Workflow: The workflow comprises three modules: (1) Phasing, where QC’d unphased cohort data is phased using SHAPEIT5 (optionally with reference panels); (2) Local Ancestry Inference, using RFMix2, GNomix or FLARE to generate ancestry segment calls from the phased VCF and reference panels; and (3) *Tractor* GWAS, which combines phased genotypes, local ancestry tracts, and covariates to perform ancestry-aware GWAS and produce ancestry-specific summary statistics. The lower panel illustrates how unphased genotypes are resolved into haplotypes, followed by local ancestry labeling using reference populations (e.g., AFR, EUR), which are then used in *Tractor* GWAS.

Accompanying this workflow, we have developed a comprehensive tutorial for all steps involved in running *Tractor* (https://atkinson-lab.github.io/Tractor-tutorial/). The tutorial explains each step from both theoretical and practical perspectives, helping users gain a deeper understanding of the *Tractor* workflow. Additionally, it provides toy data that allows users to complete a full walkthrough of the pipeline, offering hands-on experience and insight into the processes occurring behind the scenes. This tutorial lowers the barrier to entry for new users, empowering them to adopt the method in their own work and demystifying each step with this fully guided training.

High accuracy of LAI is critical, as *Tractor* relies on these estimates to perform its ancestry-based regression. One key component for accurate LAI is identifying a well-curated reference population with the ancestries present within the cohort (Honorato-Mauer *et al*., 2025). As such, we provide several pre-curated reference panels which can be utilized as part of our workflow for admixture types commonly observed in empirical data. Specifically, we have curated a two-way reference panel for samples containing predominantly African-like (AFR) and European-like (EUR) continental genetic ancestry as well as a three-way reference panel additionally containing largely homogeneous Amerindigenous-like (AMR) references, leveraging the recently released Thousand Genomes Project - Human Genome Diversity Panel (TGP-HGDP) joint-call dataset. We have benchmarked this reference panel (as compared to several alternate options for sample composition) through simulations to confirm that it provided the optimal performance for populations with a demographic model reflective of many self-identified African American individuals (Gravel *et al*., 2011).

To demonstrate the scalability and utility of our *Tractor* Nextflow workflow, we applied it in parallel to 6,245 two-way AFR-EUR admixed individuals identified from the UK Biobank, analyzing 9,957,312 variants across 32 blood biomarkers (Table S2). The workflow ran efficiently and successfully identified several established loci. For example, the trait serum ApoB identified the canonical *APOE* and *CELSR2* loci, as well as several novel, biologically plausible associations on *PCSK9* and *PMCBP1* driven by AFR ancestry (Fig. 2). The lead variants for these were absent from European ancestry tracts, highlighting the value of *Tractor* in uncovering ancestry-enriched genetic architecture.

**Fig. 2.**
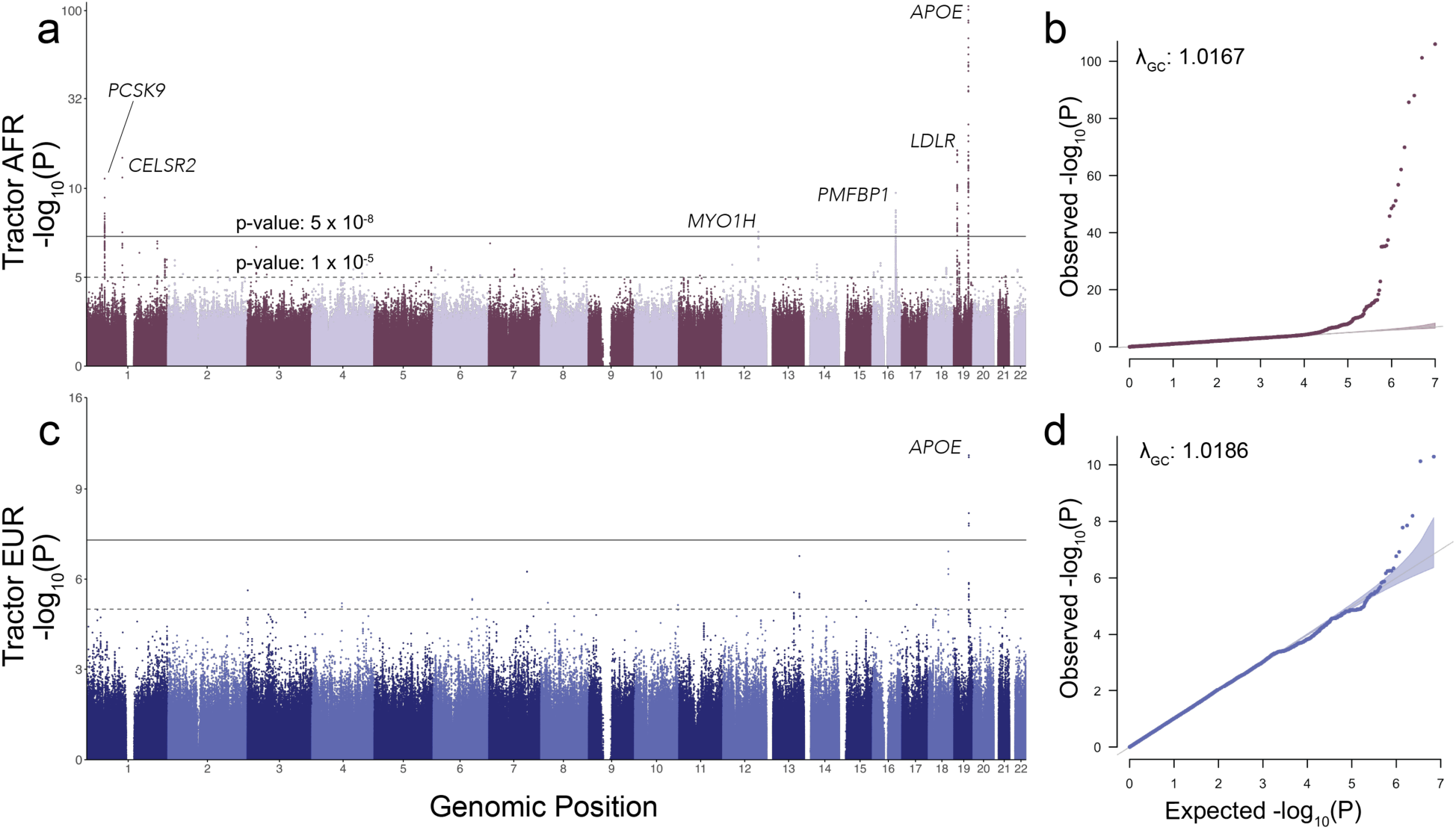
Manhattan and Q-Q plots from *Tractor* GWAS of Apolipoprotein B (ApoB) levels for AFR and EUR ancestral tracts in an admixed AFR-EUR cohort from the UK Biobank (N = 5,795). (a, c) Manhattan plots showing p-values for AFR and EUR tracts, respectively. A shared genome-wide significant locus was identified at *APOE*, with five additional significant loci detected on AFR tracts at *PCSK9*, *CELSR2*, *MYO1H*, *PMFBP1*, and *LDLR*. (b, d) Q-Q plots for AFR and EUR tracts, indicating well-calibrated Type I error control in both ancestry-specific GWAS.

We hope that the development of *Tractor* Nextflow lowers the barrier for researchers to perform association studies in admixed populations. Nextflow’s interoperability across a range of computing environments ensures that researchers can access *Tractor*’s capabilities regardless of their level of computational expertise and will lead to harmonized analyses. The scalable application of our workflow to admixed cohorts has the potential to uncover valuable genetic insights into disease architecture, identifying novel ancestry-enriched loci that may have previously been overlooked and advancing our understanding of disease biology across populations.

## Methods

### 1 *Tractor* Workflow Overview and Implementation

#### 1.1 Overview of the *Tractor* Workflow

The *Tractor* Nextflow Workflow streamlines the processing of input cohort and sequentially executes phasing, LAI, and *Tractor* GWAS. The workflow generates intermediate and final outputs from each stage, including ancestry-specific summary statistics. The pipeline is designed to scale to any number of ancestries.

This workflow’s division into independent modules allows users the option to customize the pipeline to meet their needs. The Nextflow workflow, along with its comprehensive documentation is publicly available on GitHub: https://github.com/Atkinson-Lab/TractorWorkflow.

#### 1.2 Input Requirements and Pre-Processing

The *Tractor* workflow requires input data that has undergone rigorous sample and variant-level quality control to ensure the integrity of downstream analyses, such as that recommended by the Ricopili guidelines (Lam *et al*., 2020). It is additionally typically recommended that variants be filtered for a common Minor Allele Frequency (MAF) threshold, typically 0.5-1%, and that only biallelic SNPs are retained. In addition, input VCF files should include relevant variant-level annotations, such as allele count (AC), allele number (AN), as they are required by the SHAPEIT5 software which performs the phasing (Hofmeister *et al*., 2023). Input genotypic (VCF) files can be either SNP-array or WGS data.

#### 1.3 Running the *Tractor* Workflow

##### 1.3.1 Full Pipeline Execution

The *Tractor* Nextflow workflow can be executed end-to-end with a single command, requiring only the minimal mandatory input arguments. In Module 1: Phasing, the pipeline uses SHAPEIT5 to perform genomic phasing, a gold-standard tool for biobank-scale data and is selected for its proven computational efficiency and high-quality haplotype estimation. The workflow employs the `phase_common` and `phase_ligate` tools of SHAPEIT5 to phase common variants, with support for chunked phasing and automatic ligation of these segments as two separate steps.

The workflow supports several optional arguments, including the use of a reference panel for phasing. When provided, the workflow automatically performs reference-based phasing, which is known to improve accuracy, especially when the input cohort is small (Loh *et al*., 2016). However, this might restrict the output to variants shared between input and reference VCF files.

The phased VCF serves as input to Module 2: LAI, which currently supports three widely used tools: RFMix2 (Maples *et al*., 2013), GNomix (Hilmarsson *et al*., 2021) and FLARE (Browning *et al*., 2023). These tools require a well-curated reference panel with known continental ancestry classifications and produce ancestry-inferred assignments in a commonly used format. The workflow supports configuration of multiple optional parameters for this step, and we strongly encourage users to familiarize themselves with the LAI tool settings to ensure the most accurate ancestry estimates. An example LAI plot for one of the 2-way admixed individuals from the UK Biobank is provided in Fig. S1b.

Finally, these ancestry estimates along with the phased VCF are used for Module 3: *Tractor* GWAS which consists of two steps. First, the `extract_tracts` step generates the required dosages and haplotype count files. These files, along with the phenotypic information provided by the user, are then used in the final step to perform the *Tractor* GWAS analyses in R. Linear regression is performed using the `lm()` function and logistic regression is implemented using the `glm()` function with `family` `=` `binomial(link = "logit").`

1.3.2 Running Independent Modules

The modular structure of the *Tractor* workflow allows the users to customize the execution according to their needs. For instance, users with pre-phased data or precomputed local ancestry estimates may begin their runs directly from Module 2 or Module 3, respectively. Each module can be executed independently, enabling users to integrate external tools not currently supported by this Nextflow workflow. This modularity provides both flexibility and ease of use, accommodating a wide range of input configurations and analytical preferences.

#### 1.4 Output Files and Summary Statistics

Each step in the *Tractor* Nextflow workflow produces organized output, with results stored across five main directories: `1_chunks_phased`, `2_chunks_ligated`, `3_lai`, `4_extract_tracts`, and `5_run_tractor`. When running only specific modules, only the corresponding output directories will be generated. All standard output and log files from tools executed within the workflow are stored in their respective output directories.

Module 1 (Phasing): This step performs haplotype-based phasing for common variants in chunks using `phase_common` from SHAPEIT5, with the resulting phased BCF files stored in `1_chunks_phased`. Users can choose to phase the data in chunks (recommended for large datasets) or phase it as a single block. In both cases, the final phased BCF files will reside in `1_chunks_phased`. If chunking is performed, the chunks are subsequently ligated and converted to VCF format, with the outputs saved in `2_chunks_ligated`. The phase quality score (phaseQ), which quantifies agreement in overlapping regions between adjacent chunks, is recorded in the logs file generated by SHAPEIT5. If no chunking is used, the BCF file is simply converted to VCF and stored in `2_chunks_ligated`.

Module 2 (LAI): This module performs ancestry estimation using a user-preferred tool. The resulting outputs are stored in the `3_lai` directory, with file types varying according to the selected software. RFMix2 produces local ancestry estimates (.msp file), global ancestry estimates (.Q file), as well as additional files not utilized by the *Tractor* pipeline such marginal probabilities of ancestry (.fb file) and other intermediate files. GNomix generates local ancestry estimates and marginal probabilities, as well as two subdirectories, `generated_data`, which contains the simulated data, and `models`, which contains the trained model that can be reused for LAI. The configuration file used for GNomix is also copied into the output directory to document run parameters. FLARE appends ancestry estimates directly to the input genotypic data, producing a VCF file with embedded ancestry information, and optionally includes probability estimates. A model file is also produced describing FLARE modelling parameters.

Module 3 (*Tractor*): This module consists of two steps: (1) extracting ancestral dosages and haplotype counts, which are saved as intermediate files in 4_extract_tracts, and (2) running the *Tractor* GWAS, which uses these intermediate files from previous steps and a user-provided phenotype file to generate final association statistics and is stored in 5_run_tractor.

Note that Nextflow creates multiple temporary working directories during execution and typically symbolically links final output files into the designated output directories. This can be modified in the workflow code. Most files in this workflow will be symbolically linked; however, summary statistics from the final *Tractor* step will be moved into `5_run_tractor`.

The latest release of *Tractor* provides ancestry-specific allele frequencies, local ancestry proportions, effect sizes, standard errors, t-values, and p-values for each ancestry, along with effect sizes and p-values for local ancestry terms. Additionally, the updated version supports multi-threading, enabling greater efficiency and reduced runtime.

### 2 Reference Files and Panels

#### 2.1 Reference Panel Preparation and liftOver to GRCh37

A high-quality reference panel is essential for accurate phasing and LAI, as it provides a comprehensive set of phased haplotypes that can inform the haplotype structure of study cohorts. For this purpose, we utilized the recently released joint-call dataset from the Thousand Genomes Project (TGP) and Human Genome Diversity Panel (HGDP), which expands upon the traditional TGP-only reference by incorporating individuals from a broader range of global ancestries (Koenig *et al*., 2024). This dataset is currently available only in the GRCh38 genome build.

Given that many genetic databases and resources remain aligned to GRCh37, we performed a LiftOver of the joint-call dataset to GRCh37. Initially, we filtered the GRCh38 dataset to retain only biallelic SNPs with a MAF of at least 0.005, excluding INDELs and multiallelic variants. We have also annotated each variant with allele count (AC), allele number (AN), and MAF.

Conversion to GRCh37 coordinates was performed using the LiftoverVcf tool from GATK Picard v3.1.1 (McKenna *et al*., 2010). Variants that did not successfully lift over or mapped to a different chromosome were excluded. Following coordinate conversion, the dataset was re-phased using SHAPEIT5 in a reference-free mode to generate a high-quality GRCh37-aligned reference panel. Overall, more than 99% of the variants were successfully lifted over and retained in the final dataset (Table S1).

#### 2.2 Reference Panel for Phasing

For most datasets, especially for those whose sample ancestries align with the populations represented in the TGP-HGDP joint-call dataset, this resource provides a robust reference for phasing (Koenig *et al*., 2024). For cohorts whose ancestries are absent from this panel, a custom reference may yield better results. Use of reference panels for smaller datasets has been shown to improving phasing accuracy and hence is strongly recommended (Loh *et al*., 2016). However, when using SHAPEIT5 for reference-based phasing, the resulting variant calls will be restricted to sites shared between the input cohort and the reference panel, and thus users should recommend carefully evaluating and selecting their reference strategy that maximizes phasing accuracy for their specific use case.

#### 2.3 Reference Panel for Local Ancestry Inference (LAI)

Accurate LAI requires a well-curated reference panel that matches the ancestral backgrounds of the study cohort. To support analyses of populations commonly represented in GWAS studies, we assembled reference panels from the TGP-HGDP dataset for two-way AFR-EUR and three-way AFR-AMR-EUR populations.

For the two-way AFR-EUR reference panel, we selected individuals with homogenous ancestries, restricting to continental and unrelated African and European populations. The resulting panel includes 607 African and 664 European individuals. For the three-way AFR-AMR-EUR panel, many of the AMR reference samples within the TGP-HGDP dataset are admixed. To address this, we provide a non-admixed list, containing unrelated 91 AMR individuals with over 80% AMR global ancestry as estimated by ADMIXTURE software (Alexander *et al*., 2009), along with the AFR and EUR samples described above. Users who wish to retain admixture in their AMR reference set may opt to include all available AMR samples, provided the chosen LAI software supports admixed reference panels, such as RFMix2. In such cases, we recommend adding an expectation-maximization (EM) iteration during LAI and benchmarking performance to be certain of its accuracy. All pre-curated reference individual lists are available on GitHub here: https://github.com/Atkinson-Lab/TractorWorkflow/tree/main/resources/lai_reference_panels and uploaded as Supplementary Data.

### 3 Empirical Application on 2-way AFR-EUR UK Biobank cohort

#### 3.1 Population Selection / Cohort Selection

To identify African-European admixed individuals, we utilized the Pan-UKBB classification (Karczewski *et al*., 2024), which for its own efforts has already assigned ancestry based on genetic clustering using principal component analysis (PCA), guided by reference panels from TGP and HGDP (Fig. S1a). The genetically inferred AFR category includes both homogenous AFR individuals and those with two-way AFR-EUR admixture (Fig. S1c)

As our study was conducted under a separate UK Biobank application (Application ID: 95179), it was necessary to match the sample IDs from the Pan-UKBB resource (Application ID: 31063) with those approved under our application. The Pan-UKBB dataset included 6,636 genetically inferred, unrelated AFR individuals. After bridging the datasets, we identified 6,245 individuals present in our application. This final cohort serves as the basis for all analyses presented.

#### 3.2 Pre-GWAS Quality Control

##### 3.2.1 Sample and Variant QC

For our selected cohort of 6,245 individuals, we applied stringent quality control measures to retain only high-confidence variants. Individuals with high autosomal genotype missingness (>10%) were excluded. Using the UK Biobank array-based imputed dataset, we filtered for biallelic SNPs with an imputation quality score (INFO) ≥ 0.8, a MAF ≥ 0.5%, and excluded variants with high sample missingness (>10%). All input VCF files were annotated with allele count (AC), allele number (AN), and MAF. Following filtering, 9,957,312 variants were retained for downstream analyses.

##### 3.2.2 Phenotype QC and Covariates

To demonstrate the scalability of our workflow, we analyzed all 32 blood biomarkers (Table S2) available from the UK Biobank in the selected cohort of 6,245 individuals. Since all biomarkers are continuous traits, we applied inverse rank-normal transformation to each phenotype using the RNOmni package in R, ensuring comparability and improved adherence to normality assumptions in downstream association testing.

Covariates included age at recruitment (Data Field: 21022), sex (Data Field: 31), global proportion of African ancestry as estimated by ADMIXTURE (Fig. S1c), and blood dilution factor (Data Field: 30897).

#### 3.3 *Tractor* Workflow Runs

##### 3.3.1 Arguments provided: Mandatory and Optional

The *Tractor* Nextflow workflow was executed at scale in a modular fashion. To improve phasing and LAI accuracy, we incorporated joint phasing with a reference panel. Specifically, we combined unrelated AFR (607 individuals) and EUR (664 individuals) samples from the TGP-HGDP joint-call dataset with our input cohort of 6,245 UK Biobank individuals at shared variant sites. This merged dataset was processed through Module 1 (Phasing), which employs SHAPEIT5 (v5.1.1) with a reference VCF. All chromosomes were phased in full without chunking. Because the reference panel was merged with the input cohort, non-reference-based phasing was performed. ADMIXTURE was also employed to assess global ancestry proportions.

Following phasing, the dataset was split into two groups: the original 6,245 UKBB individuals and the 1,271 AFR + EUR reference samples. The latter served as the reference panel for Module 2 (Local Ancestry Inference) with RFMix2. We ran RFMix2 with default parameters, except for the inclusion of `--reanalyze_reference true` and `--em_iterations 1`, which correspond to RFMix2’s settings to account for heterogeneity and admixture in reference populations. The global ancestry estimates obtained from ADMIXTURE and RFMix2 demonstrated a strong concordance (r² > 0.99), indicating that the LAI performed reliably in this population (Fig. S2).

Module 3, *Tractor* GWAS was run using the local ancestry estimates from the previous step, normalized blood biomarker phenotypes, and covariates using linear regression. Since the final step of *Tractor* supports multi-threading, it’s important to balance the number of variants processed (using the `--chunksize` argument) with the memory and threads provided. In our case, we used 4 CPU with 32Gb memory and a `--chunksize` of 40,000. Users are encouraged to optimize these parameters for their system, as larger chunk sizes or more threads may require increased memory.

##### 3.3.2 Computational Framework

All analyses were performed on our local HPC Cluster running Red Hat Enterprise Linux 7.9 and managed by the SLURM workload manager. Nextflow handled job orchestration, parallelization, error handling, and resource allocation across the cluster.

##### 3.3.3 Configuration File for Nextflow Workflow

All Nextflow runs require a configuration file, which defines the parameters, computational resources, and environment settings necessary to execute the workflow. This file can be tailored to support various HPC systems, job schedulers, or cloud platforms. We encourage users to consult the Nextflow documentation to explore the range of available configuration options, including specifications for CPUs, memory, file paths, and custom parameters.

Each of the *Tractor* Nextflow steps can be configured independently in terms of CPU count, memory allocation, and wall-time limits. Additionally, steps can be automatically retried with adjusted resources in the event of a failure, allowing for dynamic and adaptive execution across diverse computing environments. We allocated increased memory for the phasing and LAI steps, as these are particularly computationally intensive. For steps that failed during initial runs, the pipeline dynamically re-executed them with higher memory and CPU allocations to ensure successful completion.

## Results

### Section 1: Scalability and Computational Performance

To evaluate the scalability and robustness of our Nextflow-based workflow, we applied it to genotype and phenotype data from 6,245 individuals across 32 blood biomarkers (Table S2). Modules 1 and 2, along with the dosage extraction portion of Module 3 need to be performed only once per dataset and can be reused for *Tractor* GWAS across all traits. Only the final *Tractor* GWAS step in Module 3 needs to be executed separately for each phenotype. From a practical perspective, this means users only need to run the full pipeline once to generate the intermediate `extract_tracts` data, which represents the bulk of the compute time (∼7,500 CPU-hours). After that, running additional phenotypes requires only ∼5 CPU-hours per trait, as the workflow reuses previously generated intermediate files. For typical use, processing a new trait takes approximately 1 to 1.5 hours using 4 CPUs.

Given the scale of the analysis, we leveraged parallel execution across our local HPC cluster using configuration settings detailed in Table 1. The entire workflow required a total of 11,375 CPU-hours, with Module 1 taking ∼2,600 CPU-hours, Module 2 taking ∼4,600 CPU-hours and Module 3 taking ∼4,100 CPU-hours for all 32 blood phenotypes.

**Table 1:** Summary of computational resources used for each module of the *Tractor* workflow. The pipeline consists of three main modules and for each, we report the number of computational jobs executed and the total CPU-hours consumed. For Module 1 (Phasing), Module 2 (LAI) and Extract Tracts step of Module 3, 22 jobs were run, one for each chromosome on the genotypic data. For the *Tractor* GWAS step (Run *Tractor*), which performed linear regression separately for each of the 32 phenotypes across 22 chromosomes, a total of 704 jobs were run. As expected, phasing (using SHAPEIT5) and LAI (using RFMix2) took the largest share of computational time. We also offer median memory and CPU usage per job, as the workflow employed dynamic resource management, i.e. failed jobs were automatically rescheduled with increased resource allocations.

| Modules | Steps | Split Level | No. of Jobs | Total Wall-Clock Time | Total CPU-Hours | Median Memory/Job** | Median CPUs/Job** |
| --- | --- | --- | --- | --- | --- | --- | --- |
| Module 1 | Phasing | 22 chr | 22 | 16-12:58:04 | 105-05:26:00 | 16 Gb | 4 |
|  | Ligation | 22 chr | 22 | 0-04:14:25 | 0-04:14:25 | 4 Gb | 1 |
| Module 2 | RFMix2 | 22 chr | 22 | 12-06:13:08 | 196-03:30:08 | 64 Gb | 16 |
| Module 3 | Extract Tracts | 22 chr | 22 | 5-13:18:26 | 5-13:18:26 | 10 Gb | 1 |
|  | Run *Tractor* * | 32 phenotypes x 22 chr | 704 | 41-15:21:02 | 166-20:32:04 | 20 Gb | 4 |
| Total |  |  | 792 | 76-04:05:05 | 473-23:01:03 |  |  |

### Section 2: Application of Tractor Nextflow workflow to Apolipoprotein B levels uncovers shared and unique genetic associations

To confirm its ease of deployment, we ran the Tractor Nextflow workflow at scale across 32 traits in the UK Biobank. Here we highlight one of them, Apolipoprotein B (ApoB), as an example of the value of ancestry-specific association testing. Using the Tractor GWAS framework, we examined ApoB blood biomarker levels in 5,795 admixed individuals, leveraging local ancestry tracts to generate ancestry-specific summary statistics for African (AFR) and European (EUR) haplotypes. The average global proportions of ancestry in these samples were 92.7% AFR and 7.3% EUR (Fig. S1C). We observed well-calibrated genomic inflation factors in both ancestry tracts (AFR λ_GC_: 1.0167, EUR λ_GC_: 1.0186), showing *Tractor* well controlled for type I error (Fig. 2).

Our analysis identified six genome-wide significant loci in AFR tracts and a single hit in EUR tracts (which overlapped one of the AFR signals). This distribution is expected as the relative global proportion is higher for AFR ancestry, which confers greater power to detect significant associations in AFR tracts.

The strongest association in both ancestries localized to the *APOE* locus on chr 19. Here the lead variants differed by ancestry, but their effect sizes were very similar (β_AFR_ = – 0.66; β_EUR_ = –0.63), supporting a likely shared biological mechanism (Atkinson *et al*., 2021). The p-value differences across tracts can be linked to differences in power for AFR and EUR components. Additionally, we identified a significant AFR-specific association in *CELSR2*, consistent with previous large-scale GWAS findings related to ApoB (Richardson *et al*., 2022), further demonstrating *Tractor*’s ability to detect known biology with admixed populations.

*Tractor* also enabled the discovery of novel ancestry-specific signals. A variant near the *LDLR* locus achieved genome-wide significance in AFR tracts. While this variant has not been previously tied to ApoB directly, it has been previously associated with LDL-cholesterol and Lipoprotein(a) levels. Given *LDLR*’s role in lipid uptake, this novel locus seems to be biologically plausible. *Tractor* also identified unique hits in *PCSK9* and *PMFBP1*, both previously associated with low-density lipoprotein and total cholesterol (Wojcik *et al*., 2019). None of these variants were observed in EUR tracts (Table 2) which can be attributed to the variant’s extremely low MAF of 0.0003 in Non-Finnish European populations (gnomAD v3.1.2). These findings highlight both the power of *Tractor* to uncover ancestry-enriched loci and the broader importance of advantages of including admixed populations in genome-wide studies.

**Table 2:**
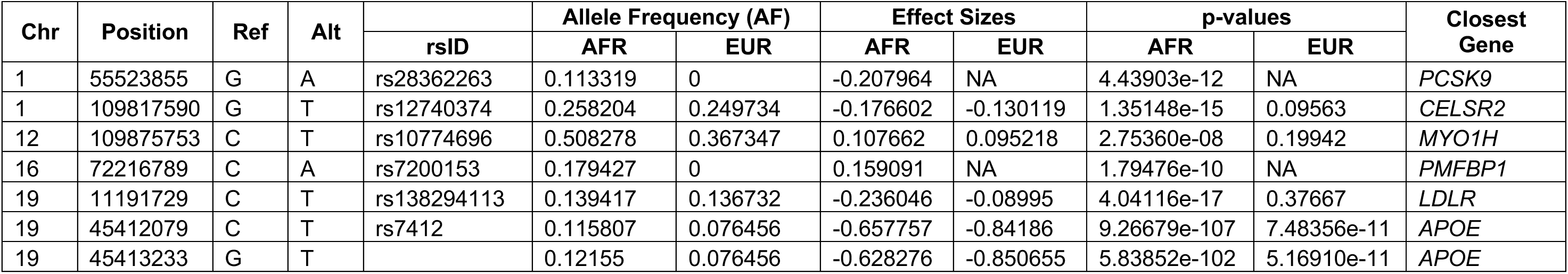
Top Genome-Wide significant loci from *Tractor* GWAS of Apolipoprotein B (ApoB) levels in Admixed AFR-EUR individuals from UK Biobank. Variants near *PCSK9* and *PMFBP1* were identified exclusively in AFR tracts due to their absence in EUR tracts (AF = 0), demonstrating *Tractor*’s ability to uncover ancestry-specific signals missed by conventional GWAS. Consequently, no effect sizes or p-values were computed for these variants in EUR tracts. The *APOE* locus showed significant associations in both ancestries but with different lead variants.

### Section 3: Tutorial

We also provide a tutorial that walks users through each step of the *Tractor* pipeline, mirroring the sequence of the Nextflow workflow. The tutorial offers a basic framework for implementing the *Tractor* pipeline and is designed to help users understand the underlying steps of the *Tractor* workflow and gain insight into how *Tractor* operates, providing transparency into its implementation beyond the automated Nextflow pipeline. The tutorial covers: 1) Phasing and LAI; 2) Optionally, correction of switch errors in phased data for 2-way admixed data; 3) Extraction of dosage and local ancestry information; 4) *Tractor* regression analysis using either Hail or a command-line tool. Complementing the Nextflow pipeline, the tutorial also includes a template for running regression analysis with Hail, a Spark-based tool that enables parallel computing that users may find the Hail implementation useful for some biobank-scale workspace environments.

We have prepared a tractable toy dataset consisting of 61 2-way AFR-EUR admixed individuals from the ASW (African Ancestry in Southwest US) subpopulation of the TGP and approximately 210,000 variants from chr 22. Using this dataset, we have simulated both binary and continuous phenotypes, assuming that only one variant within the AFR tract contributes to the trait. We also provide a reference panel for phasing and LAI, composed of 196 YRI (Yoruba in Ibadan, Nigeria) and GBR (British in England and Scotland) samples from TGP. Due to the lightweight nature of the toy dataset, users can complete the pipeline in ∼10 minutes on a typical laptop.

## Discussion

The inclusion of a broad range of populations in GWAS is crucial for several reasons (Atkinson *et al*., 2022). Firstly, exclusion of certain populations directly impacts the accuracy and applicability of genetic research for these groups. Secondly, including more representative samples aids gene discovery and better characterizes the genetic underpinnings of disease for people of all ancestry backgrounds. Per portability of GWAS findings, exclusion can miss key variants with elevated importance for understudied populations, as genetic variants and combinations of variants might be specific to certain populations, leading to incomplete understandings of diseases (The Thousand Genomes Project Consoritum, 2012; Bergström *et al*., 2020). Further, the frequency of genetic variants and patterns of linkage disequilibrium can vary significantly across populations, affecting the statistical power and robustness of GWAS findings, potentially increasing type I and II errors without proper correction.

While *Tractor* GWAS facilitates study of mixed collections and cohorts, the technical complexity and prerequisite steps required to implement *Tractor*, such as phasing and LAI pose barriers to its widespread adoption. *Tractor* Nextflow overcomes this issue by providing a user-friendly, scalable, and modular workflow that lowers the technical threshold for applying *Tractor*, helping to democratize its use in the study of admixed populations. This Nextflow workflow has a modular design allowing researchers to substitute alternatives or run individual modules independently, offering flexibility to tailor the workflow to specific datasets or computing environments. The standardized implementation also enhances reproducibility across research labs, simplifying cross-study harmonization and enabling more robust meta- and mega-analyses spanning multiple cohorts. Importantly, the workflow is also computationally efficient: while the initial steps of phasing and LAI can be time-intensive, these are performed only once on the genotypic data per cohort, enabling rapid rerunning of the final association step for additional phenotypes. The workflow is also highly parallelizable, further reducing runtime and enabling scalable application to multi-phenotype studies.

To ensure accuracy and reproducibility, we have also identified and validated optimal reference panels for both two-way (AFR-EUR) and three-way (AFR-EUR-AMR) admixed populations using the recently released joint-called TGP-HGDP dataset. This publicly available resource provides broad population coverage, allowing researchers worldwide to use a publicly available dataset that has been optimized for estimating local ancestries.

Here, we empirically demonstrate the value of this workflow by applying *Tractor* successfully to 32 blood biomarkers in 6,245 AFR-EUR admixed individuals from the UK Biobank. This large-scale effort ran in an efficient timeline and produced impactful empirical results: we successfully replicated known hits for blood lipid traits alongside identifying unique hits that were driven by ancestry-specific allele frequencies. This shows the utility of *Tractor* GWAS to find loci that may have been overlooked in previous studies using less representative cohorts or traditional GWAS methods.

Several critical considerations still remain for users. First, while *Tractor* enables ancestry-specific association testing, its statistical power depends on sufficient representation of each ancestry in the dataset. Variants that are rare within a particular ancestry or tracts that cover a small proportion of the genome are less powered for robust estimation. To maximize interpretability and power, researchers are encouraged to include ancestries with adequate sample size and sufficient genomic representation of ancestry tracts (e.g. >=5% global ancestry of the cohort). Second, *Tractor*’s performance relies on the accuracy of phasing and LAI, both of which remain active areas of methodological development, and have not been optimized for cohorts of all ancestries. Users must carefully select and optimize the references they use to get the most accurate results.

In summary, *Tractor* Nextflow provides an accessible and efficient solution for implementing ancestry-aware association testing, with the goal of expanding the inclusion of admixed individuals in genetic research. By facilitating its broader use, we hope this work will contribute to a more comprehensive understanding of the genetic architecture of complex traits across ancestries and help advance equity in genomic discovery.

## Supporting information

Supplementary Data (references)

## Acknowledgements

This research has been conducted using the UK Biobank Resource under Application Number 95179. This work was supported by funding from the National Institutes of Health. R01HG012869 has been granted to E.G.A. and R01MH106595 and R01MH134468 to C.M.N.

## Data availability

*Tractor* Workflow: https://github.com/Atkinson-Lab/TractorWorkflow

*Tractor* Workflow Documentation: https://atkinson-lab.github.io/TractorWorkflow/

*Tractor* Workflow Tutorial: https://atkinson-lab.github.io/Tractor-tutorial/

