## Supplementary File for "Tractor Workflow Pipeline: A Scalable Nextflow Framework for Local Ancestry-Aware Genome-Wide Association Studies"

### Supplementary Tables and Figures

| | LiftOver | | | | | | LiftOver (Post-Filters) | | SHAPEIT5 Phasing (filter)*** | |
|---|---|---|---|---|---|---|---|---|---|---|
| chrom | Original number of variants* | Variants lifted over | Percent Variants lifted over | Variants failed to lift over | Percent Variants failed to lift over | Variants lifted over, but with mismatched ref alleles | LiftOver Filtered Variants | Percent LiftOver Filtered Variants | Variants after SHAPEIT5 phasing (MAF filter: 0.5%) | Percent Variants after SHAPEIT5 phasing (MAF filter: 0.5%) |
| chr1 | 1,135,111 | 1,127,990 | 99.37% | 4,399 | 0.63% | 2,722 | 1,126,431 | 99.24% | 1,122,580 | 98.90% |
| chr2 | 1,208,210 | 1,206,077 | 99.82% | 1,605 | 0.18% | 528 | 1,205,894 | 99.81% | 1,202,383 | 99.52% |
| chr3 | 1,032,523 | 1,029,864 | 99.74% | 2,088 | 0.26% | 571 | 1,029,735 | 99.73% | 1,027,060 | 99.47% |
| chr4 | 1,051,797 | 1,050,900 | 99.91% | 507 | 0.09% | 390 | 1,050,880 | 99.91% | 1,047,568 | 99.60% |
| chr5 | 941,101 | 940,682 | 99.96% | 337 | 0.04% | 82 | 940,662 | 99.95% | 938,568 | 99.73% |
| chr6 | 941,665 | 934,115 | 99.20% | 6,402 | 0.80% | 1,148 | 933,839 | 99.17% | 932,035 | 98.98% |
| chr7 | 855,823 | 848,016 | 99.09% | 6,079 | 0.91% | 1,728 | 847,734 | 99.05% | 846,046 | 98.86% |
| chr8 | 805,762 | 804,219 | 99.81% | 447 | 0.19% | 1,096 | 804,200 | 99.81% | 802,237 | 99.56% |
| chr9 | 639,850 | 636,333 | 99.45% | 3,068 | 0.55% | 449 | 635,364 | 99.30% | 633,797 | 99.05% |
| chr10 | 733,631 | 725,522 | 98.89% | 7,744 | 1.11% | 365 | 725,174 | 98.85% | 722,919 | 98.54% |
| chr11 | 724,378 | 722,955 | 99.80% | 981 | 0.20% | 442 | 722,804 | 99.78% | 720,049 | 99.40% |
| chr12 | 696,024 | 692,817 | 99.54% | 3,070 | 0.46% | 137 | 692,759 | 99.53% | 690,870 | 99.26% |
| chr13 | 525,417 | 524,754 | 99.87% | 613 | 0.13% | 50 | 523,496 | 99.63% | 521,409 | 99.24% |
| chr14 | 472,985 | 470,625 | 99.50% | 756 | 0.50% | 1,604 | 470,623 | 99.50% | 469,242 | 99.21% |
| chr15 | 426,670 | 425,270 | 99.67% | 1,175 | 0.33% | 225 | 425,201 | 99.66% | 423,901 | 99.35% |
| chr16 | 454,875 | 450,948 | 99.14% | 3,903 | 0.86% | 24 | 450,658 | 99.07% | 449,262 | 98.77% |
| chr17 | 394,768 | 388,705 | 98.46% | 4,633 | 1.54% | 1,430 | 388,188 | 98.33% | 387,257 | 98.10% |
| chr18 | 415,068 | 414,880 | 99.95% | 128 | 0.05% | 60 | 414,857 | 99.95% | 413,357 | 99.59% |
| chr19 | 327,725 | 325,617 | 99.36% | 692 | 0.64% | 1,416 | 325,612 | 99.36% | 325,085 | 99.19% |
| chr20 | 324,182 | 319,625 | 98.59% | 4,350 | 1.41% | 207 | 319,261 | 98.48% | 318,579 | 98.27% |
| chr21 | 201,577 | 200,118 | 99.28% | 1,361 | 0.72% | 98 | 199,148 | 98.80% | 198,695 | 98.57% |
| chr22 | 204,523 | 198,019 | 96.82% | 5,168 | 3.18% | 1,336 | 194,093 | 94.90% | 193,774 | 94.74% |
| **Total** | **14,513,665** | **14,438,051** | **99.48%** | **59,506** | **0.41%** | | **14,426,613** | **99.40%** | **14,386,673** | **99.13%** |

**Table S1: Variant summary statistics filtering, liftOver and phasing of the TGP-HGDP Joint Dataset from GRCh37 to GRCh37.** The original dataset, released in GRCh38, was first filtered to retain only biallelic SNPs with MAF > 0.5%, forming the baseline set reported in the "*Original number of variants*" column. All subsequent percentages are relative to this set. A total of 99.48% of variants were successfully lifted over to GRCh37, while 0.41% failed to map. To ensure positional accuracy, we retained only variants that mapped to the same chromosome as their original location, yielding 99.40% in the “LiftOver (Post-Filters)” set. This filtered dataset was then phased using SHAPEIT5 (`phase_common`) with a consistent MAF filter, resulting in a final retention of 99.13% of the original variants. This high-quality GRCh37-converted reference set was used for downstream phasing and local ancestry inference.

| # | UKBB Phenocode | Phenotype Description | Total Cases |
|---|---|---|---|
| 1 | 30000 | White blood cell (leukocyte) count | 5564 |
| 2 | 30060 | Mean corpuscular haemoglobin concentration | 5564 |
| 3 | 30600 | Albumin | 5418 |
| 4 | 30610 | Alkaline phosphatase | 5840 |
| 5 | 30620 | Alanine aminotransferase | 5838 |
| 6 | 30630 | Apolipoprotein A | 5390 |
| 7 | 30640 | Apolipoprotein B | 5795 |
| 8 | 30650 | Aspartate aminotransferase | 5806 |
| 9 | 30660 | Direct bilirubin | 4865 |
| 10 | 30670 | Urea | 5837 |
| 11 | 30680 | Calcium | 5416 |
| 12 | 30690 | Cholesterol | 5836 |
| 13 | 30700 | Creatinine | 5836 |
| 14 | 30710 | C-reactive protein | 5827 |
| 15 | 30720 | Cystatin C | 5835 |
| 16 | 30730 | Gamma glutamyltransferase | 5835 |
| 17 | 30740 | Glucose | 5409 |
| 18 | 30750 | Glycated haemoglobin (HbA1c) | 4667 |
| 19 | 30760 | HDL cholesterol | 5412 |
| 20 | 30770 | IGF-1 | 5800 |
| 21 | 30780 | LDL direct | 5825 |
| 22 | 30790 | Lipoprotein A | 4843 |
| 23 | 30800 | Oestradiol | 1514 |
| 24 | 30810 | Phosphate | 5408 |
| 25 | 30820 | Rheumatoid factor | 441 |
| 26 | 30830 | SHBG | 5369 |
| 27 | 30840 | Total bilirubin | 5802 |
| 28 | 30850 | Testosterone | 5292 |
| 29 | 30860 | Total protein | 5417 |
| 30 | 30870 | Triglycerides | 5835 |
| 31 | 30880 | Urate | 5831 |
| 32 | 30890 | Vitamin D | 5680 |

**Table S2: Summary of Phenotypes Analyzed in Tractor GWAS of UK Biobank Admixed AFR-EUR Individuals**. List of all 32 blood-biomarkers analyzed using Tractor GWAS in an admixed AFR-EUR cohort from the UK Biobank along with its corresponding UK Biobank phenocode, description of the phenotype, and the number of individuals with available data used in the analysis.

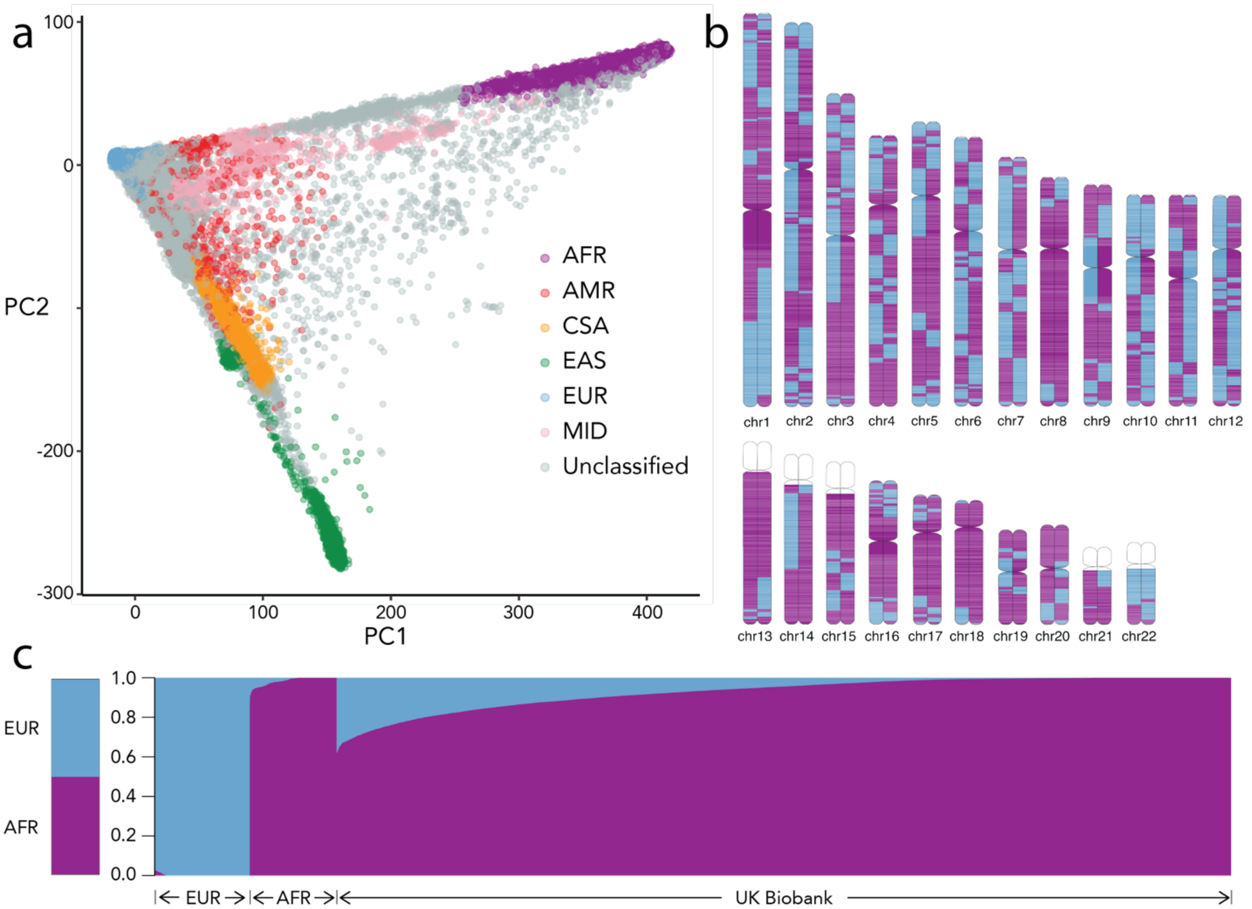

**Figure S1: Overview of cohort selection and ancestry inference for Tractor GWAS in UK Biobank.** (a) Principal Component (PCA) plot of all UK Biobank individuals. Individuals classified as AFR by Pan-UKBB[17] (in purple) were selected for inclusion in Tractor analyses. (b) Local Ancestry for one of the selected admixed individuals as inferred by RFMix2 showing segmental ancestry along the genome with alternating AFR and EUR tracts (c) ADMIXTURE plot (K=2) showing ancestry proportions for the final set of 6,245 admixed AFR-EUR individuals, alongside AFR and EUR reference samples on the left from the TGP-HGDP joint-call dataset.

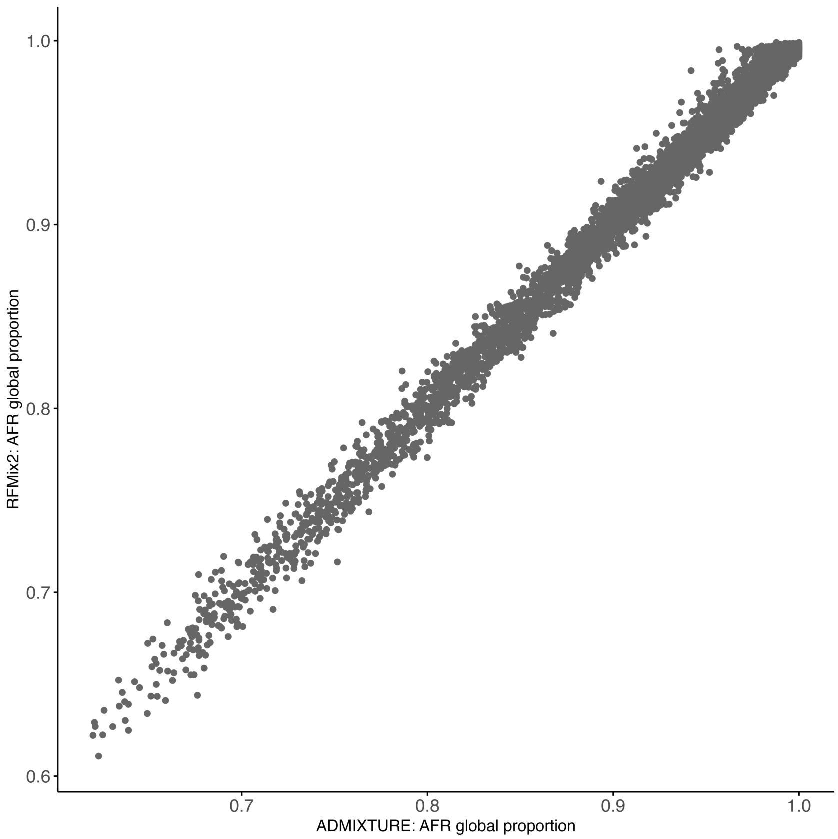

**Figure S2: Strong correlation between lobal Ancestry Estimates from ADMIXTURE and RFMix2 Supports Accuracy of Local Ancestry Inference using RFMix2.** Global ancestry proportions estimated using ADMIXTURE (K=2) were compared to global ancestry estimates from RFMix2 in 6,245 admixed AFR-EUR individuals from the UK Biobank. The two methods demonstrated strong concordance ($r^2 > 0.99$), supporting the accuracy and reliability of RFMix2's local ancestry inference in this cohort.
